# The Cyanobacterial *dnaX* Intein Encodes an Out-of-Frame Homing Endonuclease and is Sporadically Distributed in the Phylum

**DOI:** 10.64898/2026.09.14.751524

**Authors:** Daniel S. Phillips, Stella DeSimone, Johann Peter Gogarten

## Abstract

Inteins are self-splicing proteins with a wide distribution in single-celled organisms, viruses, and plastids. The function of inteins remains an unanswered question. They may be regulatory elements as they must splice out of their host protein. However, recent research findings point to inteins likely being mobile selfish genetic elements capable of horizontal transfer. Here the intein inhabiting cyanobacterial DNA Polymerase III gamma/tau subunit (*dnaX*) serves as a case study into the conservation and phylogenetics of the intein compared to the host extein. Phylogenetic analysis reveals a sporadic distribution of intein alleles having an evolutionary history that is incongruent with the extein that may reticulate independently across clades. The *dnaX* intein can also contain an out-of-frame homing endonuclease (HE), a rarity for annotated inteins, that may have been the result of secondary invasion of an HE into a mini-intein. The simultaneous presence of intein-free and intein-containing alleles in populations of *Microcystis* aeruginosa challenges current theories about the life cycles of homing endonuclease containing selfish genetic elements and suggests that alternatives to the homing cycle need to be considered for intein persistence. Our analysis reveals an incongruent, disjointed distribution of this specific intein, suggesting that it is neither conserved among individual cyanobacterial lineages nor a critical component of *dnaX* functionality yet is able to persist in populations without being lost.

## Introduction

Inteins represent an enigmatic part of genomic ecosystems. These self-splicing genetic elements (Hirata 1990; Kane et al. 1990; Perler 1997) have been found in every kingdom except in the nuclear genome of multicellular metazoans (inteins are present in the genomes of multicellular fungi (Poulter et al. 2007)) and are especially widespread in unicellular organisms (Gogarten et al. 2002), often present in conserved regions of critical housekeeping enzymes (Swithers et al. 2009). While the mechanics of splicing are understood and have been co-opted for the creation of novel proteins (Anastassov et al. 2024) among other biochemical applications (Pavankumar 2018), their actual function and purpose relative to the organism as a whole remain unresolved and is a matter of some debate (Gogarten 2002; Gogarten and Hilario 2006; Pietrokovski 1994).

After the discovery of inteins; researchers proposed that inteins were functional regulatory elements (Belfort 2017; Kelley et al. 2016, 2018; Woods et al. 2020), as the enzyme is not functional until the intein spontaneously splices out either post or co-translationally, as splicing times vary depending on the specific intein (Shah et al. 2012). Thus, inteins could serve as a blunt instrument of post translational control. This view has been supported by numerous *in vitro* assays where splicing is inhibited by irregular conditions such as temperature, pH, or the presence of heavy metal ions. Researchers hypothesize that these compounds and conditions may prevent the protein folding or otherwise interfere with amino acid residues critical to the splicing reaction. This ability has been co-opted by protein biochemists to better control protein activity by engineering an intein from a thermophile into a protein of interest of a mesophile as the intein will not splice until subjected to its native temperature (McNeal et al. 2025). However, the conditions in these experiments may affect the functionality of any number of proteins and may not reflect intein function in the living organism (Belfort 2017; Lennon et al. 2021; McNeal et al. 2025; Woods et al. 2020). In instances of genes invaded by multiple inteins, an inhibitor only affects a specific intein, while other inteins are unaffected and continue to splice out successfully; as was observed with the class III intein that invades mycobacterial *dnaB* replicative (Tori and Perler 2011; Woods et al. 2020). Regardless, multiple inteins are present in genes or in the subunits of a holozyme in a manner that would be more suggestive of a selfish genetic element (SGE) that functions evolutionarily independently of its host. Currently, relatively little focus is on the evolution of inteins, not only on their presence or absence in a genome or a clade, but also to their rates of evolution at the nucleotide level compared to their host gene.

Analysis of nucleotide substitution patterns can reveal evolutionary pressures for the maintenance of conserved regions. These approaches have been frequently used for the analysis of viral (Bush et al. 1999), bacterial and archaeal (Kogay et al. 2026) and eukaryotic genes (Saccone et al. 2025), especially in investigating selfish genes (Goddard and Burt 1999; Werren 2011) which are generally known to evolve and mutate faster than protein coding or otherwise functional genomic sequences (McLeod and Gandon 2025) as most functional genes are under greater purifying selective pressures. By examining substitution rates and gain/loss events, we can determine whether HEs are being maintained, degraded, or lost as full inteins become mini-inteins. This helps distinguish among competing models for intein persistence: does the intein invasion follow the homing cycle model with each phase going to completion (Goddard and Burt 1999) or does the intransitive fitness relationship (Barzel 2011) between the different intein alleles (e.g. an intein-free gene and an intein-containing homolog) allow for the long-term coexistence of the different intein alleles.

Prokaryotic genomes are notably susceptible to deletion pressures for non-functional genes in the absence of strong selection pressures (Mira et al. 2001). Accordingly, strong selection pressure should be exhibited on the key residues in the splicing domain (SD), which must remain as constant as the residues comprising the active site in a functional enzyme, and in the HE, so that the gene would have means so invade empty alleles, whereas the other sites would not have pressure to be conserved since the fate after splicing would be degradation and recycling. There has also been little empirical focus on the evolution and distribution of mini inteins (those lacking a HE) compared to their full-intein counterparts. Though there are many examples in public databases of these mini and full intein alleles; most of the research is either theoretical in nature focusing on the possible homing cycles of similar mobile elements like introns is currently unknown how or why and how often an intein may gain or lose a HE, and if the HE should be treated as a separate parasitic selfish genetic element or as part of the intein. One theory posits that the loss of a HE is a natural consequence of an intein invading all empty sites in a population. Once the intein is fixed in the local population, the evolutionary pressure for an intein to maintain a functional HE disappears, the HE then gradually degrades as the smaller mini-intein may incur less of a cost compared to the longer full intein. Without the means to invade empty target sites the mini-intein may then itself be lost; outcompeted by an intein-free allele and unable to reinvade empty sites.

This homing cycle model (Goddard 1999; Gogarten 2006) does not take into account that the intein or the HE might gain a function beneficial to the host organism or the host organism’s population. The latter was discussed for different SGEs (McLeod 2025). In case of inteins a benefit to the population or species could occur if inteins with HED in different genome locations are not fixed in a population. Parasexual mechanisms could bring together genomes with different HE specificities. These would produce double strand cuts in the other genome thus forcing recombination to assemble an uninterrupted complete genome. Also, the homing cycle might operate in different parts of a wider, heterogenous population, or local environmental conditions might either favor the spread of inteins or the loss of intein containing alleles. Another possibility is that perhaps some selfish genetic elements have a negligible fitness cost and therefore are not strongly selected against (Werren 2011) as may be the case with inteins that invade single copy genes such as replicative DNA polymerases since they are produced only once during the life cycle of a bacterium and in relatively small numbers compared to other key housekeeping or metabolic protein that would be continuously synthesized. Finally, the stable coexistence of inteins with HED, inteins without HED and host genes without inteins was determined to be possible even in well mixed populations (Barzel 2011; Yahara 2009).

The intein that is present in the *dnaX* gene in cyanobacteria has several qualities that make it an ideal test case to study intein evolution and dynamics. *dnaX* encodes the gamma/tau subunit of the principal replicative polymerase of cyanobacteria, DNA Polymerase III. It is invaded by a single intein in a highly conserved region of the clamp-loader domain of the protein. Other researchers have noted that this intein is present as both a full intein allele and a homologous mini-intein allele with both having very similar splicing domains. The gamma/tau subunit is not the only intein-containing component in the replisome: *dnaE* and *dnaB* are both noted to contain multiple inteins in cyanobacteria and other lineages as well (Caspi 2003; Gogarten 2002). While the *dnaX* intein has been annotated in public databases since its discovery and characterization (Liu 1997), an in-depth study of the different alleles and their structure has not yet been published, with the exception of an X-ray crystallographic study of the mini-intein in *Spirulina platensis* (Boral et al. 2020). Here we describe the cyanobacterial *dnaX* intein, the coexistence of the different *dnaX* alleles in the same populations over time, and the selection pressures acting on the intein whose functions (self-splicing and HE) are encoded by separate, overlapping reading frames.

## Methods

### Phylogenetic analysis

All sequences were obtained from NCBI and homology was determined using BLASTp or tBLASTn searches (Altschul 1990). Preference was given to homologs that came from the complete genomes of cultured isolates. *dnaX* homologs were taken from major families of Cyanobacteria (Dvořák 2025) in an attempt to capture the diversity of the phylum, from non-photosynthetic sister taxa (Moore 2019) to multicellular heterocyst-forming (Kumar 2010) and true-branching species (Gugger 2004; Koch 2017). The finalized dataset consisted of 282 *dnaX* amino acid sequences of which 105 were intein-free, 85 were mini-inteins and 92 were full-intein. The single gene trees were constructed using IQ-TREE (Nguyen 2015), with the model finder program used to determine the most probable maximum likelihood model. IQ-TREE was also used to compare artificially constrained trees to the maximum likelihood tree. Sequences were aligned using MUSCLE (Edgar 2004) in Seaview (Gouy 2021).Ultra-fast bootstrap support (UFBS) (Minh 2013) values determined using 10,000 iterations were used to assess branch support. Phylogenic trees were visualized and annotated in Figtree (Rambaut 2018). The extein tree was rooted using a homolog from *Escherichia coli* strain ST540 (GenBank: CP007265.1) a distantly related Gram-negative species which has no publicly available sequences containing this specific intein. IQTree3 run logs, parameters, and alignments are included in the supplemental data.

### Constrained Tree Topology Tests

Artificially constrained extein trees were constructed that forced each *dnaX* allele (intein-free, full-intein, mini-intein) into separate mono-clades. These artificial trees were compared to the ML extein phylogeny to determine if there is any support of monophyly with the approximately unbiased (AU) test (Shimodaira 2002) in IQTree3. To test topologies to determine congruent evolution of the full-intein, the extein, and HE domain trees were generated with those amino acid sequences and compared to the full intein tree topology using the same methods.

### Structure prediction

*DnaX* homologs from *Microcystis aeruginosa* (MA) were used to create predicted structures with Alphafold3 in ChimeraX (Pettersen 2021). ORF finder in NCBI and HHpred were used to determine the existence of a +2 ORF that encodes a presumptive functional HED. The presence of the out of frame HE was further confirmed via BLASTn and BLASTp searches that resulted in close homologs of know LAGLIDAG HE from a yeast intron (I-Wcal). While only MA predicted structures were included in this analysis, multiple structures were generated from various genomes for initial analysis.

### Mutation Rate Comparison

A set (n=130) of MA and Nostocales unique full-intein-containing DNA Polymerase III gamma subunits (the smaller subunit of *dnaX* which contains the intein) sequences were used to create alignments without any gaps (793 codons total). The dN/dS (omega) values at each position of both the *dnaX* ORF and the HE ORF (+2ORF) were determined using the FUBAR program as implemented in DataMonkey (Weaver 2018). Figures were constructed using python matplotlib and Microsoft excel.

### Metagenomic Presence of multiple alleles

Publicly available Illumina HiSeq mWGS runs were selected from NOAA Great Lakes Environmental Research Laboratory (GLERL) and Great Lakes Atlas for Multi-omics Research (GLAMR) metagenomics projects that monitor and repeatably sample specific locations in Lake Erie for harmful algae blooms (HABs) as a long-term metagenomic shotgun sequencing project. The simultaneous presence of different MA *dnaX* alleles (intein-free, mini-intein, full intein) as assessed using custom probes specific and exclusive to each specific MA *dnaX* alleles (S1). Probes were confirmed by blastn to only have significant matches to MA species present in Lake Erie’s watersheds (shown to have over 97% identity with MA sequences while other cyanobacteria shared less than 80% identity). The presence of sequences specific to these alleles was confirmed using the medium sensitivity alignment algorithm on Geneious Prime and only reads that aligned with the probe were counted.

## Results

### Predicted structure of the dnaX intein

The intein structure predicted with AlphaFold3 contained a low-confidence disordered region in all sequences tested, as opposed to the higher-confidence structures of the intein SD and extein (Fig. 1A, D). A predicted high-confidence structure that resembles a LAGLIDADG-class HE was identified in an open reading frame (ORF) that overlaps with the disordered intein sequence between the N- and C-terminal SDs (Fig. 1B). This was supported by aligning the AlphaFold3-generated structure with a known LAGLIDADG HE, which showed similarities, particularly in the antiparallel β-sheets and α-helices that comprise the sequence motifs typical of this HE class (Fig. 1C). Further blastn results showed sequence homology to known HE. The +2ORF also has putative cyanobacterial Shine-Dalgarno motifs upstream of the start codon.

**Figure 1.**
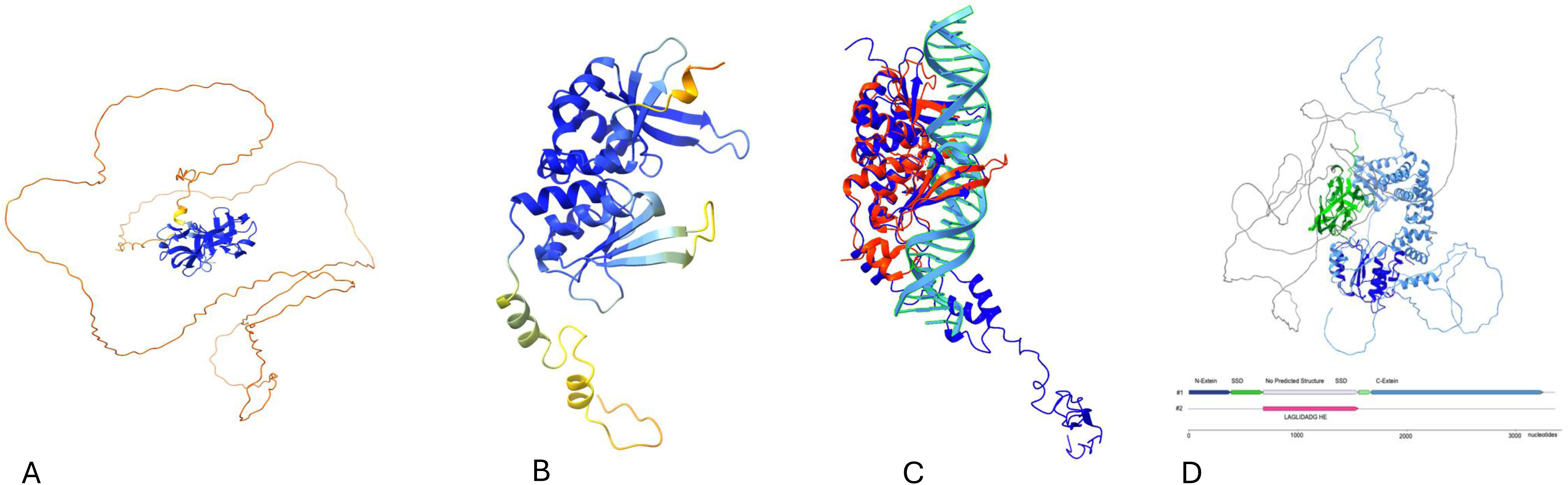
AI predicted structure of the *dnaX* intein and putative HE in the +2 ORF. (A) Intein inframe Alphafold3 prediction of a *Microcystis aeruginosa* DNA Poll III gamma/tau subunit (NCBI Reference Sequence: WP_185240695.1) showing of splicing domains with a high degree of structural confidence (in blue) and low confidence (in orange and red) where a HED was previously assumed to be. (B) shows the Alphafold3 predicted structure +2 ORF of that intermediate with a higher degree of confidence and is consistent with resolved LAGLIDAG homing endonucleases (C) Resolved intronic HED I-Wcal, PDB 6X1J, (red) overlaid with predicted HED (blue) bound to DNA (light blue) using ChimeraX matchmaking (D) AlphaFold 3 predicted structure of DNA Poll III tau subunit N and C exteins (blue) N and C intein splicing domains (green) and HED (grey) map to the gene with the alternate HED +2ORF in magenta.

### dnaX Extein and Intein Amino Acid Phylogenies and Topology Tests

The dnaX extein tree, rooted using an *Escherichia coli dnaX* homolog as an outgroup, showed that both full and mini-intein alleles are widespread among known later branching cyanobacterial clades such as *Nostocales, Dolichospermum, Planktothrix,* and *Anabaena.* Inteins were notably absent in the known earliest branching clades such as *Gloeobacter* and the non-photosynthetic sister phyla (*Melainabacteria)* and in species with notably reduced genomes such as *Prochlorococcus*. In numerous closely related clades, all three alleles are present, and this observation is supported by high ultrafast bootstrap (UFBS) values; most dramatically in *Microcystis* genomes where the exteins are almost identical at the amino acid level (Fig. 2). The AU-test results of comparing the consensus tree to artificial constrained trees show a much lower likelihood for phylogenies with imposed monophyly of full-inteins, mini-intein, or mini-inteins. This was also in accordance with other topology tests (Table 1A).

**Figure 2.**
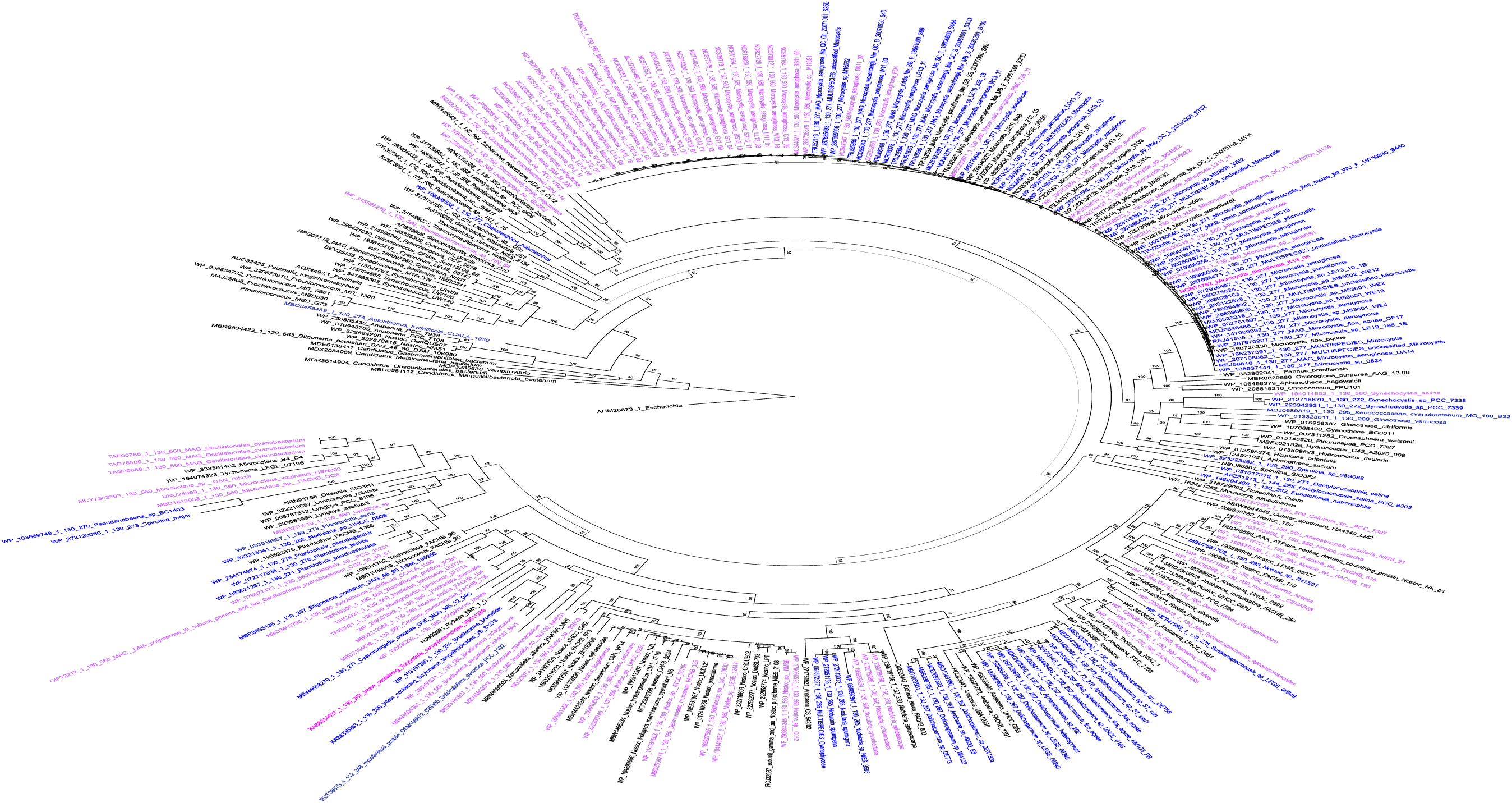
*dnaX* Extein Tree from Cyanobacteria and Sister Taxa genomes. Phylogenetic tree of the amino acid sequence of the DNA Pol III gamma/tau subunit (*dnaX*). Colored by allele type: intein-free (black), mini-intein (blue), full HED intein (purple). UFBS support indicated at each node with highly supported (>85) branches in bold. Substitution model used: JTT+F+I+R7.

**Table 1.** Topology tests comparing. (A) Extein trees constrained by intein presence to the ML constrained tree. (B) full-intein trees to the corresponding HE domain trees, extein trees, and the splicing domain trees. Topology tests were done on IQTree3.

**A AU Tests on Extein Trees**
| Tree | p-AU<sup>1</sup> | logL<sup>2</sup> | deltaL<sup>3</sup> | bp-RELL<sup>4</sup> | p-KH<sup>5</sup> | p-SH<sup>6</sup> | c-ELW<sup>7</sup> |
| --- | --- | --- | --- | --- | --- | --- | --- |
| Unconstrained Tree | 1 | -117736.1015 | 0 | 1 | 1 | 1 | 1 |
| Intein-free constrained | -1.6e-07 | -123733.3425 | 5997.2 | 0 | 0 | 0 | 0 |
| Full Intein Constrained | -3.12e-44 | -124399.2129 | 6663.1 | 0 | 0 | 0 | 0 |
| Mini Inteins Constrained | -1.16-49 | -122342.8068 | 4606.7 | 0 | 0 | 0 | 0 |

**B AU Tests Comparing Full Intein Trees to Extein Tree**
| Tree | p-AU | logL | deltaL | bp-RELL | p-KH | p-SH | c-ELW |
| --- | --- | --- | --- | --- | --- | --- | --- |
| Full Intein Tree | 0.906 | -24197 | 0 | 0.894 | 0.884 | 1 | 0.894 |
| HE Domain | 0.0939 | -24247.89352 | 51.036 | 0.106 | 0.116 | 0.467 | 0.106 |
| Extein | 2.16E-66 | -24772.15406 | 575.3 | 1 | 0 | 0 | 4.04E-155 |
| Splicing Domain | 5.10E-60 | -24742.96708 | 546.11 | 0 | 0 | 0 | 8.70E-95 |

<sup>1</sup>p-AU is the probability of the AU-test, <sup>2</sup>logL is the absolute likelihood value of the tree, <sup>3</sup>deltaL is the difference in likelihood of the tree with the highest likelihood. Bp-RELL is the proportion of resampled replicates in which the tree has the highest likelihood. <sup>4,5,6,7</sup>p-KH, p-SH, c-ELW are alternative, less conservative topology tests indicating the probability of a given tree being the true topology.

The unrooted intein phylogeny (Fig. 3) does not provide strong statistical support for monophyly of either intein-containing allele, a pattern consistent with multiple independent losses of the HE. However, monophyly of intein types within MA is strongly supported, with high UFBS support separating the mini-intein and full-intein clades. Topology tests (Table 1B) comparing the full-intein tree to the corresponding extein, SD, and HE trees showed that both the SD and extein topologies were significantly incongruent with the full-intein topology and were strongly rejected by the AU test (AU = 5.10 × 10□□□ and 2.16 × 10□□□, respectively). In contrast, the HE topology was not rejected by any test metric (AU = 0.0939, p-SH = 0.467, p-KH = 0.116) and exhibited a substantially lower ΔL value (51.0) than either the SD (546.1) or extein (575.3) trees. Together, these results indicate a greater degree of evolutionary congruence between the HE domain and the full-intein sequence than between the full intein and either the host extein or the intein splicing domain.

**Figure 3.**
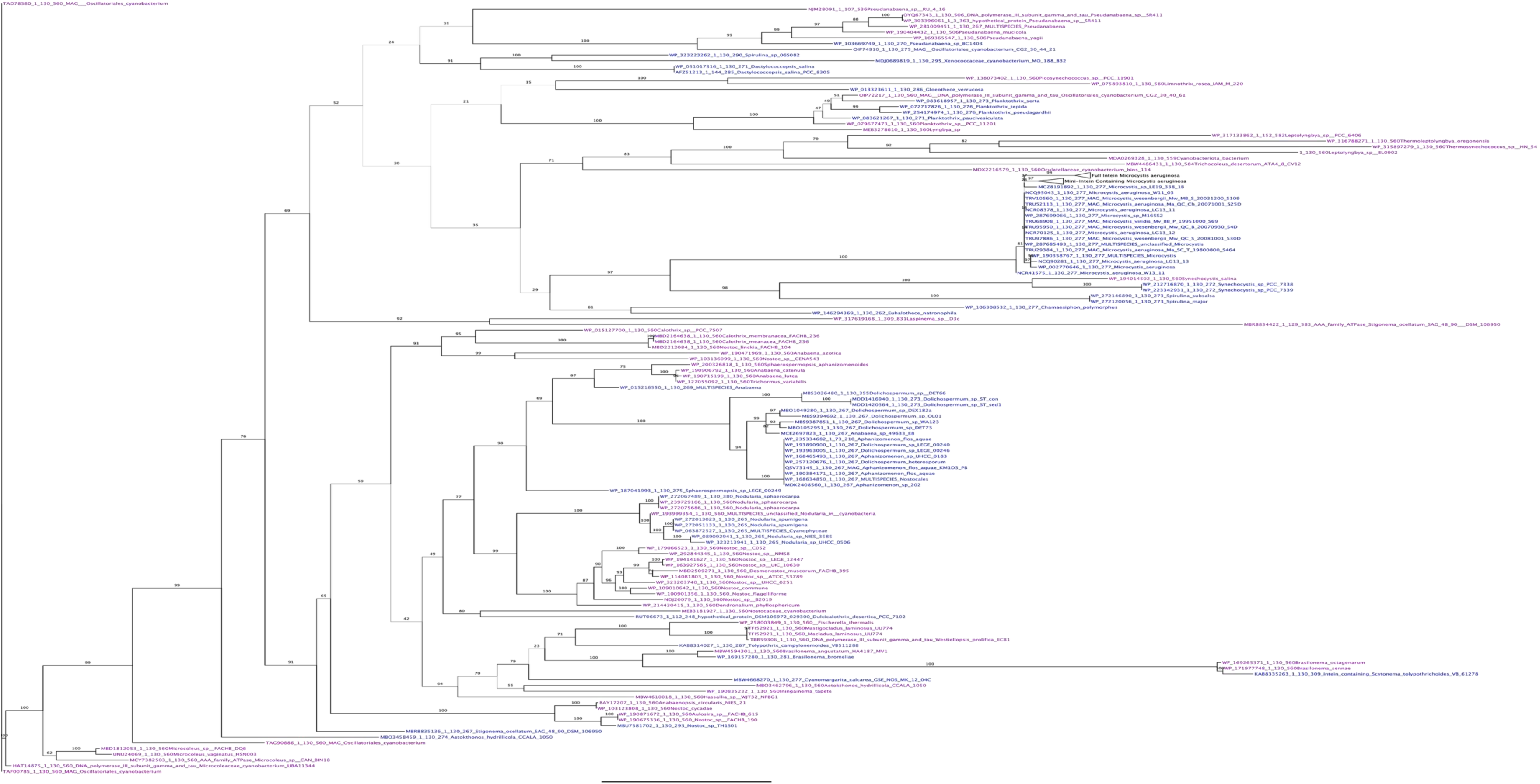
*dnaX* intein tree of representative taxa. Unrooted phylogenetic tree of the intein amino acid sequence from Cyanobacteria and sister phyla genomes. Colored by allele type: mini-intein (blue), full HED intein (purple). UFBS support indicated at each node with high supported branches (>85) in bold. Substitution model used: GTR20+F. MA monophyletic clades are collapsed for clarity.

### Co-existence of multiple alleles in a population

Metagenomic sequencing runs (Table 2) containing MA 16S rRNA reads also contained reads aligning to all three *dnaX* allele classes (intein-free, mini-intein, and full-intein) throughout the 2014-2021 sampling period. The samples were collected from adjacent locations and therefore likely represent the same regional population of planktonic MA. The continuous presence of all three allele classes in every sequencing run containing MA *dnaX* reads indicates long-term coexistence of intein-free, mini-intein, and full-intein alleles within the same geographic population.

**Table 2.**
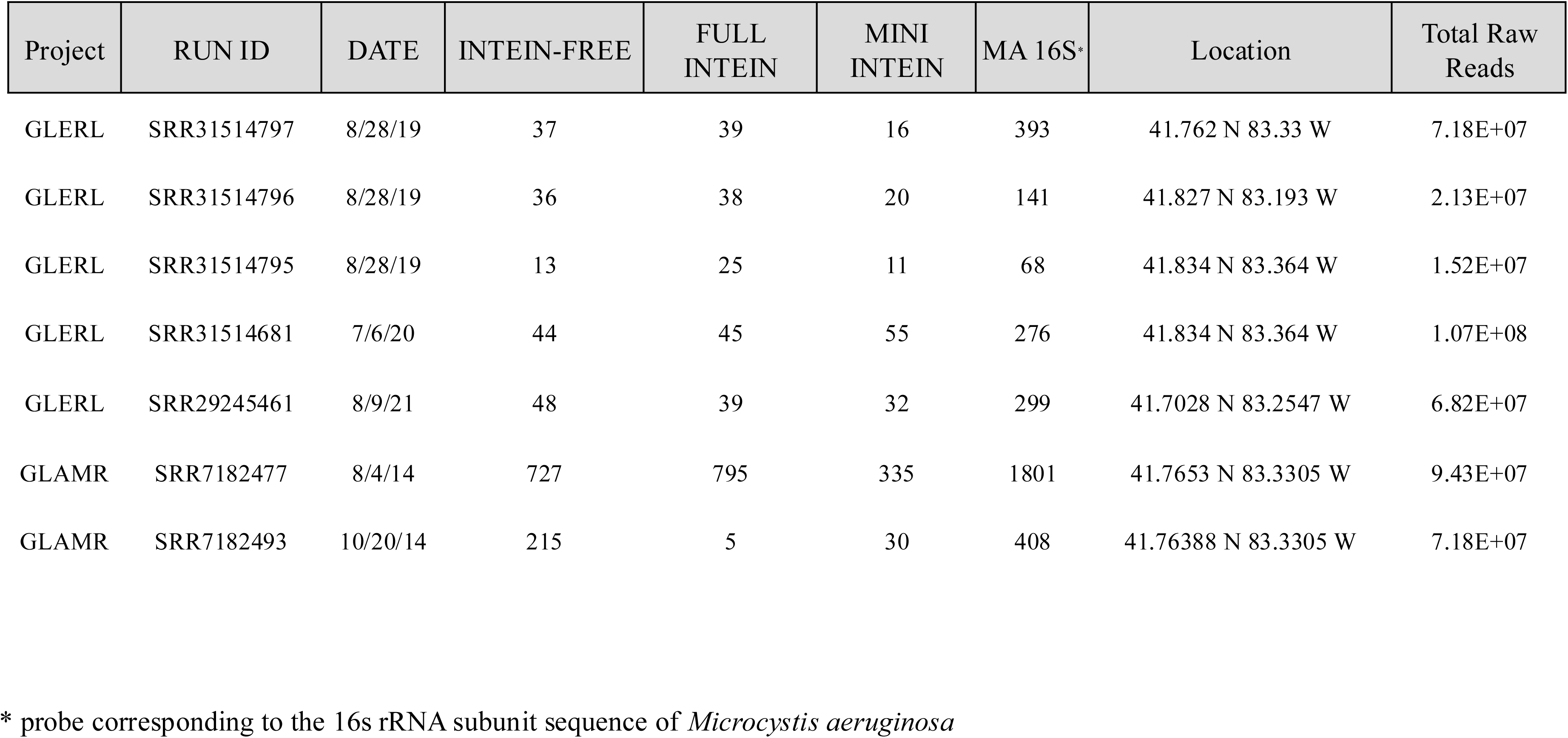
Raw read counts of each MA *dnaX* allele in Lake Erie metagenomic sequencing runs. Read count of each allele of each allele matching the to template probes from publicly available environmental monitoring projects including meta data of collection date and location. Map of template sites in supplementary figure 1.

| Project | RUN ID | DATE | INTEIN-FREE | FULL INTEIN | MINI INTEIN | MA 16S* | Location | Total Raw Reads |
| --- | --- | --- | --- | --- | --- | --- | --- | --- |
| GLERL | SRR31514797 | 8/28/19 | 37 | 39 | 16 | 393 | 41.762 N 83.33 W | 7.18E+07 |
| GLERL | SRR31514796 | 8/28/19 | 36 | 38 | 20 | 141 | 41.827 N 83.193 W | 2.13E+07 |
| GLERL | SRR31514795 | 8/28/19 | 13 | 25 | 11 | 68 | 41.834 N 83.364 W | 1.52E+07 |
| GLERL | SRR31514681 | 7/6/20 | 44 | 45 | 55 | 276 | 41.834 N 83.364 W | 1.07E+08 |
| GLERL | SRR29245461 | 8/9/21 | 48 | 39 | 32 | 299 | 41.7028 N 83.2547 W | 6.82E+07 |
| GLAMR | SRR7182477 | 8/4/14 | 727 | 795 | 335 | 1801 | 41.7653 N 83.3305 W | 9.43E+07 |
| GLAMR | SRR7182493 | 10/20/14 | 215 | 5 | 30 | 408 | 41.76388 N 83.3305 W | 7.18E+07 |

\* probe corresponding to the 16s rRNA subunit sequence of *Microcystis aeruginosa*

### Selection pressures at each site the two open reading frames

A subset of 123 unique MA, *Nostocales* and *Anabaena* full-intein containing *dnaX* nucleic acid sequences were selected to create a gap-free alignment to determine substitution biases at each codon site. The resulting omega values demonstrate frame dependent selection pressures. The extein and parts of the intein splicing domain have an omega<1. In the region of the HE however there is a dramatic shift in the number of sites where omega>1. In the +2ORF however, the omega value falls below 1 in many sites. Many sites in the +2ORF have an omega value close to that of the extein (Fig 4).

**Figure 4.**
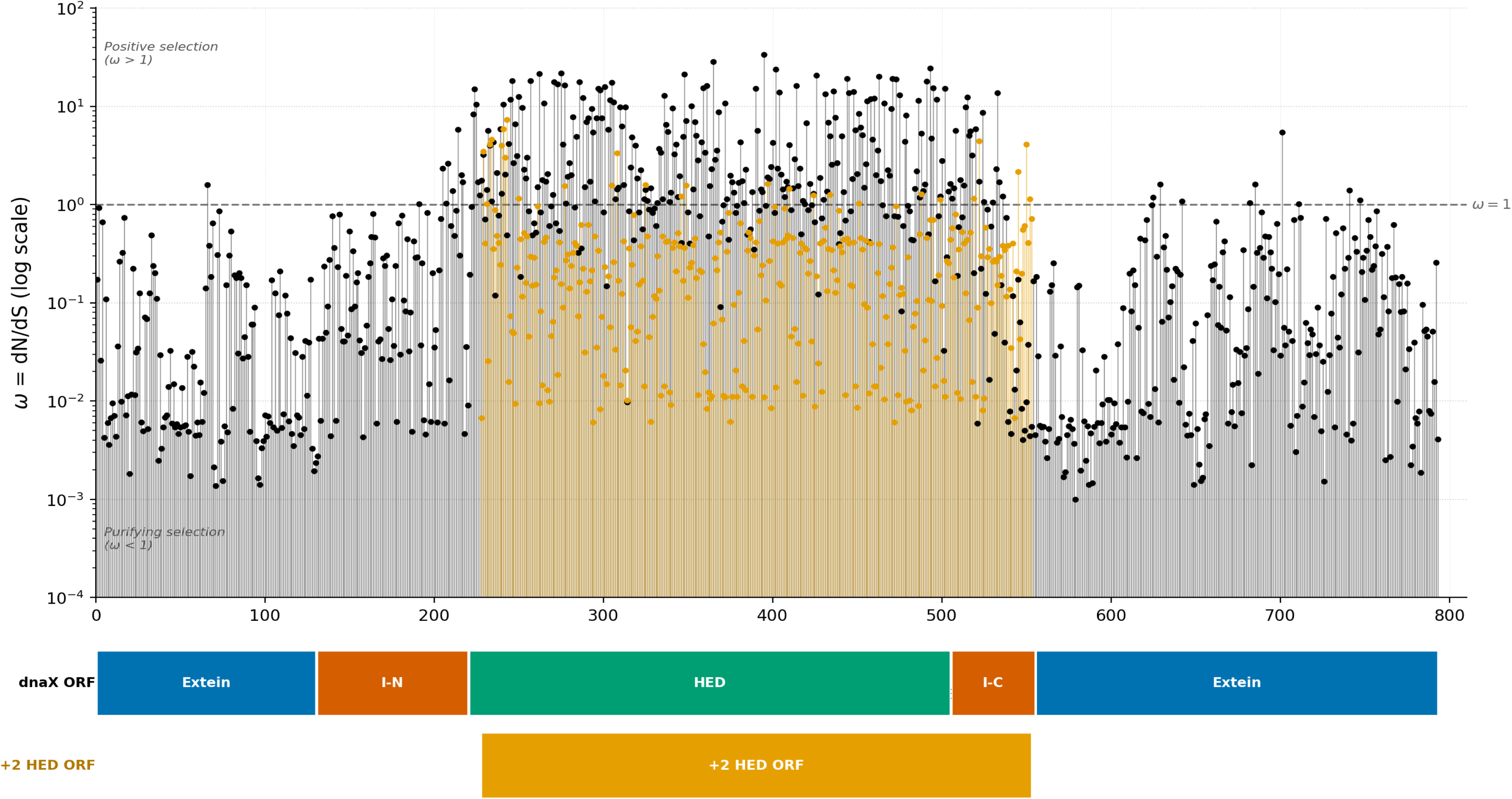
Site-wise comparison of Omega values for the *dnaX* ORF and the +2 ORF HE. The omega value at each site in an aligned subset (n=123) of unique full-intein containing MA and *Nostocales* and *Anabeana* genes. Analysis was run on the alternate +2 ORF (gold) of the putative HED (green). I-N and I-C (orange) are the intein splicing domains and the extein (blue) is the gamma/tau subunit itself. Analysis was done using FUBAR in DataMonkey

## Discussion

### Phylogenetic evidence for intein gain and loss

The hypothesis of inteins being conserved regulatory elements is underpinned by *in vitro* experiments that demonstrate the inhibition of protein splicing by external environmental factors and thereby preventing the extein from becoming functional. Inteins, like any enzyme, are sensitive to insults such as oxidative stress or presence of reactive metal species that interferes with the internal interactions of the splicing residues, especially in non-canonical or class III splicing mechanisms (Kelley et al. 2018). Those experiments mainly focus on *in vitro* assays (Panda et al. 2021) so the actual function in a natural system remains unresolved; is splicing inhibition a side effect, or a feature? Due to a lack of conclusive *in vivo* evidence, a bioinformatics approach determining the level of within lineage conservation may provide some illumination. The *dnaX* intein, due to being widespread in both full-intein and mini-intein forms in cyanobacteria provides an opportunity to further investigate the level of conservation, the selection pressures, and population dynamics to help determine if this intein is functioning as a selfish genetic element, a regulatory element, or provides another function selected at the organismal or group level. It would be expected that if inteins were providing a function relative to the host gene its presence would be conserved in lineages (Ågren 2016) and the intein’s evolution would largely tract with that of the extein. A selfish gene however would exhibit multiple instances of gain and loss, as nonfunctional genes are prone to loss in prokaryotic genomes (Ochman 2009; Mira et al. 2001).

The phylogeny modeled here paints a picture that is inconsistent with strict vertical inheritance. The intein is absent from many known early-branching cyanobacteria including their non-photosynthetic sister taxa: *Melainabacteria* and *Obscurales* (Tan et al. 2024). Notable intein-free clades include *Prochlorococcus* genomes and its (geologically) recent ancestor that became and endosymbiont of *Paulinella* (Reyes-Prieto et al. 2010), along with *Synechoccus* and *Pseudanabaena* genomes. These intein-free early-branching taxa could indicate that the intein is not a basal feature of the core cyanobacterial genome and instead may be the result of a later addition (or invasion). In many known later-branching taxa (Boden et al. 2025) such as *Nostocales*, *Anabeana*, *Dolichospermum*, and *Fischerella* both the mini and full intein are present in representative genomes. These taxa are noted for their comparably larger genomes, and heterocystic metabolic and complex structural phenotype and as opposed to earlier branching taxa which are known to have comparatively reduced genomes (Zhang et al. 2024), and may have more complex regulatory pathways (Moya et al. 2020). However, throughout the phylogeny there are numerous examples of recent homologs with intein gain and loss events present in highly supported branches; indicating that the loss of an intein is not a great detriment to the organism; an observation further supported by prior *in vitro* fitness experiments (Naor et al. 2016). Furthermore, AU-tests comparing the topologies of the consensus tree to artificially constrained trees for the both the mini and full intein show that the constrained trees are completely rejected. While the phylogeny was created using sequences intended to capture major cyanobacteria families it is by no means exhaustive and it is possible that taxa not yet sequenced will reveal a wider distribution of the *dnaX* intein. Additionally, some genomes are represented by MAGs, that may still be contaminated despite screening, rather than sequenced from isolated cultures.

The intein phylogeny mirrors the patchy distribution observed across the cyanobacterial species tree. Full inteins and mini-inteins are interspersed throughout the phylogeny while generally clustering by taxonomic group. This pattern is particularly evident in *Nodularia*, *Pseudanabaena*, and *Planktothrix*, where mini-intein splicing domains are often nearly identical in amino acid sequence to those of full inteins. The close similarity between these sequences suggests that the HE has been gained and lost repeatedly throughout evolutionary history rather than being removed in a small number of ancient events. One notable exception is the highly supported branch separating MA mini-inteins from MA full inteins (Fig. 3). Given that the corresponding splicing domains differ by only a small number of amino acid substitutions, this separation was unexpected. Resolution of deeper relationships within the intein phylogeny remains limited, as bootstrap support decreases toward the base of the tree. This is consistent with previous observations that inteins may evolve more rapidly than their host genes, making ancient evolutionary relationships difficult to resolve and leaving the timing of both intein and HE invasion events uncertain in this analysis.

Additional support for reticular gain and loss events across clades comes from topology tests comparing the unconstrained maximum-likelihood phylogeny with alternative constrained topologies. Trees in which intein-free exteins, mini-inteins, or full inteins were forced into separate monophyletic groups were strongly rejected by AU and related topology tests (Table 1A). If the intein had been inherited primarily through vertical descent following a limited number of gain or loss events, these constrained topologies would be expected to provide a reasonable fit to the data. Their rejection instead supports a history of repeated intein gain and loss throughout cyanobacterial evolution and is more consistent with the behavior of a mobile genetic element than a stably inherited component of the dnaX locus. The topology tests comparing the full-intein, SD, HE, and extein (Table 1B) also reject congruency of the full intein and the extein topology. The inability to reject the HE topology supports the full intein acting as single unit rather than the HE acting a separate gene, given its presence in an alternate ORF. The major anomaly of this analysis is the strong rejection of the SD topology compared to the HE. This may be an artifact due to the smaller number of sequences in the alignment over all combined with the fact that several residues are identical or at least highly conserved across the alignment due to their critical role in splicing. This leads to a comparably smaller percentage of phylogenetically informative sites that may be the root cause of the discordance.

### A functional HE is encoded in an overlapping reading frame

To demonstrate the +2ORF encodes a functioning HE, a substitution rate analysis was conducted on a subset of full-intein sequences from MA and *Nostocales*. Restricting the analysis to closely related taxa allowed the construction of high-quality nucleotide sequence alignments with zero gaps while maximizing the power to detect selection. All sequences were unique at the nucleic acid level so that the analysis would not be biased due to genetically identical homologs. The FUBAR model (citation) was selected because this Bayesian approach is well equipped for fast analysis of large datasets with unconstrained selection parameters among codon sites. This is particularly relevant for inteins, where functionally essential splicing residues are highly conserved while other regions have been shown to evolve substantially faster than their host exteins (Pietrokovski, 2001; Butler et al., 2006; Arsenault et al., 2025).

Most of the resulting dN/dS values are broadly consistent with patterns reported for other inteins. The extein exhibited a low dN/dS ratio suggestive of purifying selection, consistent with the highly conserved function of DNA Pol III subunits encoded by *dnaX*. In contrast, much of the intein sequence displayed elevated dN/dS values, suggesting reduced evolutionary constraint outside of the residues directly involved in protein splicing. The +2ORF with the putative HE showed the opposite pattern with low omega values. In particular, residues associated with the conserved LAGLIDADG motifs (codons 450 and 500) were subject to strong purifying selection, suggesting that maintenance of HE function may be evolutionarily important (Fig. 4). This frame-dependent level of selection generates higher dN/dS values in the first ORF since a synonymous substitution in this ORF is likely to be non-synonymous. This increased dN/dS ratio in the overlapping region of the extein ORF (Fig. 4) is not due to an increase in non-synonymous substitutions (dN) but rather reflects the dramatic drop in the number synonymous substitutions in the extein reading frame in the overlap region (Supplementary Fig. 2).

One explanation for these contrasting patterns is that the two components are subject to different selective pressures. Within the *dnaX* reading frame, the intein may experience relatively weak constraint provided that splicing activity is maintained. The HED, however, would be expected to remain functional to facilitate homing and the continued spread of the intein through unoccupied alleles. If selection in the acts primarily to preserve endonuclease activity, the overlapping portion of the extein frame is free to accumulate mutations as long as these are synonymous in the HE frame. This interpretation is consistent with the view that the *dnaX* intein has not evolved an essential host-associated function and instead persists through the activity of its homing endonuclease, preventing displacement by intein-free alleles.

The first description of the *dnaX* intein assumed that the HE was encoded in-frame with the remainder of the intein (Liu 1997), although homologous mini-inteins lacking the HE were also noted. Later analysis would group *dnaX* with other non-canonical inteins when it was proposed that HE homologs were found in alternate open readings within the *dnaX* intein splicing domains based on homology to known phage HE sequences particularly double DNA binding motifs in the HE which were spuriously assigned in the initial discovery paper (Gorbalenya 1998). An actual structure of the HE which would further validate the dnaX intein being non-canonical remained unresolved, either through either X-ray crystallography or AI models, until this present analysis. As there are few documented examples of broadly distributed inteins containing an out-of-frame HE, this arrangement may represent a secondary invasion event rather than a simple frameshift-derived modification of an ancestral full intein. Such a scenario could help explain both the rarity of this architecture in other intein systems and its widespread occurrence among cyanobacterial *dnaX* inteins (Novikova 2016). While AI-modeling was not done exhaustively on all sequences, BLAST databases searches and the lack of in-frame HE in any alignments suggests that this frame-shifted HE is the result of an insertion event, although subsequent blast searches have been unable to determine what that exact ancestor HE would have come from, either a phage or other endogenous HE.

The continued presence of mini-inteins in cyanobacterial clades and in MA populations seems counterintuitive to current proposed theories of the homing cycle, where it is presumed to be an evolutionary dead-end for the selfish genetic element (Gogarten 2006). Plausible scenarios for the persistence of a mini-intein may include an increased amount conjugation or other HGT events that allow these genes to persist absence a pressure to maintain them. The MA metagenomes suggest an alternative; all three alleles may co-exist with other cyanobacterial populations. Deeper sequencing depth may reveal this pattern in more populations, and this only became evident in this study due to the significant focus on MA populations and its high level of genetic similarity in a given ecotype (Cai 2003) which makes it simple to discern MA reads from other cyanobacterial strains. Speculations for these observations include: the simultaneous presence may only be passing, needing a larger time scale for the intein containing alleles to be lost in a population, or perhaps on a population level maintaining a certain percentage of intein-containing alleles facilitates genetic recombination and evolution which would be a long-term benefit to maintaining genetic diversity in the absence of sexual reproduction. Further genetic analysis on single nucleotide polymorphisms and synteny between bacteria in intein-containing and intein-free populations would be needed to further examine possible increased rates of genetic recombination from populations with widespread inteins.

## Conclusion

The phylogenies of individual inteins are often neglected, yet they do provide a useful framework for determining whether these elements play a functional role relative to the host organism or instead behave as selfish genetic elements. In the case of the cyanobacterial *dnaX* intein, the phylogeny reveals multiple independent gain and loss events, rejection of topologies consistent with strict vertical inheritance, and complete absence from several cyanobacterial lineages, suggesting that the intein is not a necessary component of the core cyanobacterial genome and that closely related taxa are largely tolerant to its absence. The intein also contains an unusual out-of-frame HED, strongly supported through protein modeling, that is highly conserved and subject to purifying selection, indicating that maintenance of a functional homing endonuclease remains important for the persistence and spread of full intein alleles. AU topology tests further suggest that the HED and splicing domain largely share the same evolutionary history and provide little evidence that the HED behaves as a fully independent genetic element. In MA populations from Lake Erie intein-free, mini-intein, and full-intein alleles coexist over multiple years, a pattern that is difficult to reconcile with simple homing-cycle models that predict eventual fixation of HED-containing alleles. If similar patterns occur in other cyanobacterial populations, it may indicate that intein-containing and intein-free alleles coexist for longer evolutionary periods than currently appreciated and raise the possibility that inteins provide a benefit at the population level rather than the genomic level, potentially through increased opportunities for recombination and the maintenance of genetic diversity.

## Authorship Credit

JPG and DP designed the study, wrote, and edited the manuscript. DP conducted the phylogentics, structural and metagenomics analysis. SD conducted the dN/dS analysis.

## Acknowledgements

We thank Danielle Arsenault, Sophia Gosselin, and Yutian Feng for knowledgeable support throughout the course of this project as well the MCB department at UConn for funding through summer fellowships, and the University of Connecticut’s Computational Biology Core at the Institute for Systems Genomics for providing computational resources.

---

**Supplementary Figure 1.**
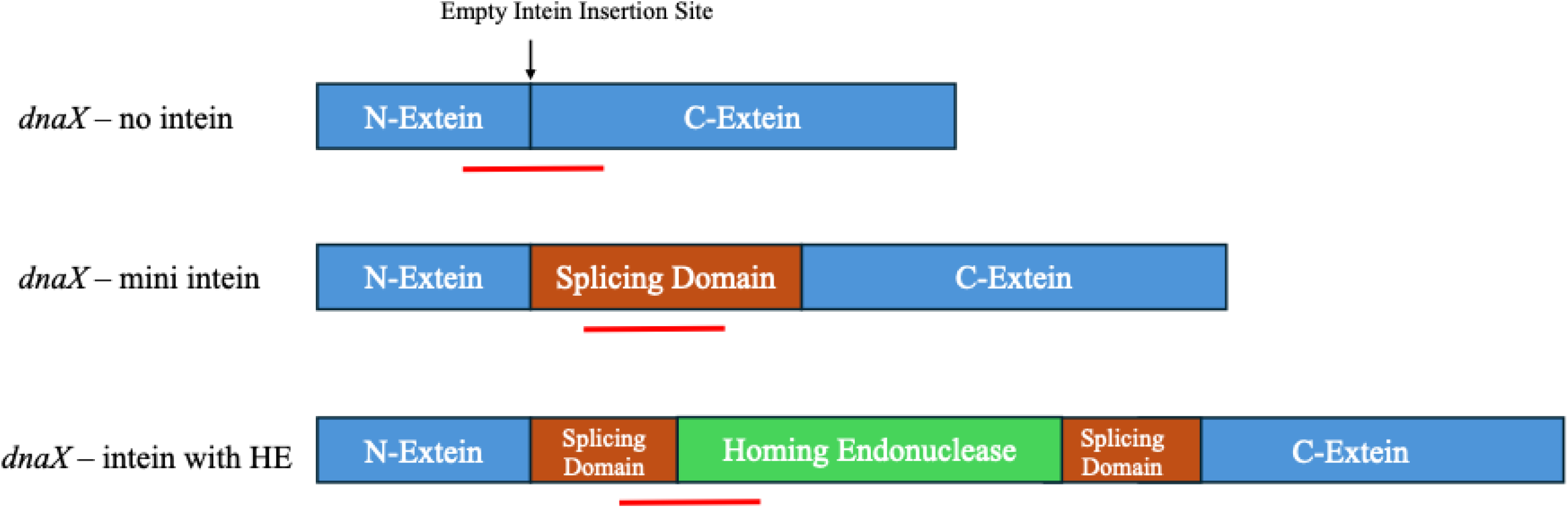
Diagram of allele specific probes to MA *dnaX* alleles in metagenomes. Diagrams shows the binding sites of the probes (red line) which are specific to a single *dnaX* allele and exclusionary to the other two. Only reads that covered the majority of the probe were included in analysis. Reads were taken from public ally available freshwater metagenomes sequenced on Illumina platforms.

**Supplementary Figure 2.**
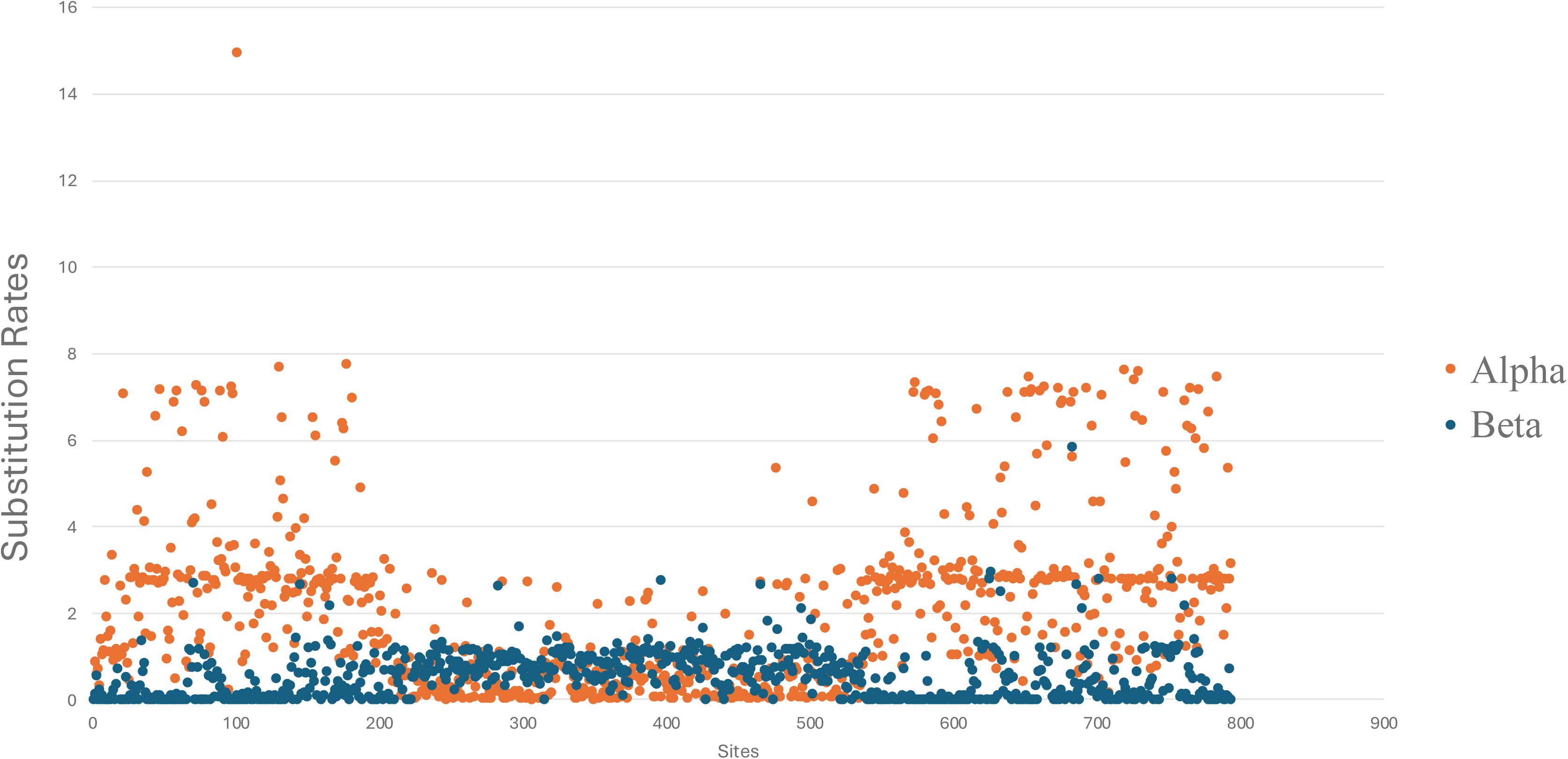
Alpha and Beta Rates *dnaX* Full Intein containing subset. Substitution rates at each amino acid position the extein ORF of a subset of Nostoc and MA *dnaX* homologs calculated in DataMonkey FUBAR. Alpha (Orange) represents dS or synonymous substitutions and Beta (Blue) represents dN or non-synonymous substitutions

